# Wnt signalling controls abscission dynamics in mouse embryonic stem cells

**DOI:** 10.64898/2026.03.05.709800

**Authors:** Snježana Kodba, Beatriz Morales Lasierra, Erika Timmers, Agathe Chaigne

## Abstract

Cell division is a crucial process that ensures proper development of multicellular organisms. Cell division ends in abscission, a process in which the intercellular bridge between two sister cells is cut. Although abscission usually happens shortly after chromosome segregation, abscission is severely delayed in mouse embryos and embryonic stem cells (mESC). The regulation of the duration of abscission influences cell fate transitions but how cell state and abscission dynamics crosstalk remains unknown. Here, we show that a key pluripotency pathway, the Wnt signalling pathway, controls abscission dynamics. Upon deactivation of Wnt signalling in naïve mESCs, abscission becomes faster. Wnt signalling regulates abscission dynamics in mESCs through two mechanisms. First, Wnt signalling keeps the amount of Aurora B high at the intercellular bridge, probably by preventing Aurora B degradation. In turn, high Aurora B activity at the bridge delays bridge abscission. Second, a key component of Wnt signalling, the kinase GSK-3β localizes at the intercellular bridge with microtubules and their associated proteins (MAPs). In pluripotent cells, inactivation of GSK-3β leads to an increase of stable microtubules at the bridge stable which causes delayed abscission. Crucially, inhibition of GSK-3β after cells have exited naïve pluripotency accelerates abscission, demonstrating that cell state influences the output of the abscission signalling pathway. The permissive function of canonical Wnt on cell state is thought to be mediated by reinforcement of existing pluripotency network; altogether, our work shows that non-canonical Wnt is also context dependent.

## Introduction

During cell division, the genetic and cytoplasmic content of a cell get separated into two daughter cells. This is orchestrated by the mitotic spindle, a multicomponent, microtubule-rich machinery. During the separation of the two sister chromatids, the central spindle is formed between the two daughter cells (White and Glotzer, 2012). The central spindle trigger the constriction of an actomyosin ring which leads to the formation of an intercellular bridge connecting the two daughter cells (Mierzwa and Gerlich, 2014). Intercellular bridges are microtubule-rich membrane tubes containing the midbody (Kodba and Chaigne, 2023a). The division ends in abscission, a membrane scission event mediated by the ESCRT-III complex which finally separates the two sister cells (Carlton and Martin-Serrano, 2007; Elia et al., 2011; Guizetti et al., 2011).

In most mammalian cells, the intercellular bridge is cut 1-2 hours after its formation (Gershony et al., 2014). However, there are many exceptions (Kodba and Chaigne, 2023a). In pluripotent stem cells the abscission is delayed for up to 12 hours (Chaigne et al., 2020; Kodba and Chaigne, 2023). When mESCs exit naïve pluripotency, abscission becomes faster, and the intercellular bridge is cut in 2-3 hours (Chaigne et al., 2020). We have previously shown that the timing of abscission in mESCs depends on the stability of the microtubules in the bridge under the dependence of the activity of Aurora B kinase (Kodba et al., 2025). Naïve mESCs, which undergo slow abscission, display high levels of Aurora B and stable microtubules at the bridge during bridge maturation, while cells that have exited naïve pluripotency have less Aurora B, less stable microtubules, and faster abscission. Aurora B activity is responsible for microtubule stability, though whether this is direct or indirect remains unknown. In turn, stable microtubules delay abscission by preventing MCAK recruitment to the midbody (Kodba et al., 2025).

While pluripotency controls abscission dynamics through regulation of Aurora B activity and microtubule stability, we do not understand how pluripotency maintaining signals control Aurora B and microtubules. In mESC, pluripotency is maintained through inputs integrated at various levels (Kinoshita and Smith, 2018a). Several signalling pathways regulate the pluripotency of mESCs, including Wnt signalling (Huang et al., 2015). Interestingly, proteins of Wnt signalling have also been shown to regulate microtubule stability and cell division in various cell types (Bufe et al., 2021; Ciani et al., 2004; Darmasaputra et al., 2024; Fumoto et al., 2012), pinpointing the Wnt pathway as a possible candidate for abscission regulation across cell states.

Wnt signalling is an evolutionarily conserved pathway that orchestrates fundamental biological processes, including the determination of cell fate, tissue morphogenesis during development, and the maintenance of adult tissue homeostasis (Reya and Clevers, 2005; Steinhart and Angers, 2018). There are two primary branches of Wnt signalling: the canonical pathway (also known as Wnt/LRP5/6 or β-catenin-dependent) and the non-canonical pathway (β-catenin-independent) (Xue et al., 2025). Canonical Wnt signalling is initiated by the simultaneous binding of Wnt ligand to the Frizzled (Fzd) and LRP5/6 receptors, which recruits Dishevelled (Dvl) (Gammons et al., 2016; Kang et al., 2022). This induces LRP5/6 phosphorylation by Glycogen Synthase Kinase-3β (GSK-3β) and Casein Kinase 1γ (CK1γ) and the recruitment of the “destruction complex” (Cselenyi et al., 2008) containing notably the GSK-3β and CK1α. kinases. When canonical Wnt is inactive, GSK-3β and CK1α phosphorylate β-Catenin. This phosphorylation marks β-Catenin for ubiquitination, which ultimately leads to its degradation by the proteasome (Li et al., 2012; Ranes et al., 2021). In embryonic stem cells, Wnt signalling is activated, and the stabilized nuclear β-catenin acts as a transcriptional regulator that removes TCF3-mediated repression of pluripotency factors and supports naïve pluripotency (Fan et al., 2020; Faunes et al., 2013; Kelly et al., 2011).

Non-canonical Wnt pathways are also activated by the binding of Fzd/LRP5/6 receptors to Wnt ligands and their activity also depends on GSK-3β kinase (Qin et al., 2024). In particular, the Wnt-Dependent Stabilization of Proteins (Wnt/STOP) (Acebron et al., 2014) is active during cell division. When Wnt/STOP is inactive, GSK-3β can phosphorylate other proteins besides β-Catenin, which leads to their degradation (Acebron and Niehrs, 2016). However, when Wnt/STOP is active, GSK-3β is inactive and its targets then accumulate. This protein accumulation regulates cell size during cell division suggesting that Wnt/STOP pathway is important to counteract protein degradation in dividing cells with reduced transcription and translation activity (Acebron et al., 2014). The Wnt/STOP pathway stabilizes around 15% of cellular proteins, probably due to the relative promiscuity of the consensus sequence and priming sequence of GSK-3β (Sutherland, 2011a), including proteins involved in DNA remodelling, cell-cycle progression, and regulation of the cytoskeleton,(Stolz and Bastians, 2015). Finally, the kinase GSK-3β, sits at the intersection of signalling pathways and has hundreds of known substrates. In particular, GSK-3β can phosphorylate several microtubule binding proteins, such as CLASP1/2 (Kumar et al., 2009; Watanabe et al., 2009), and modulate their function on microtubule dynamics(Aher et al., 2020; Grimaldi et al., 2014; Lawrence et al., 2023).

Here, we explore the role of Wnt signalling on the duration of abscission, using mESCs as a pluripotent model system. Together, our results demonstrate that Wnt signalling is a major regulator of abscission. We show that active Wnt signalling in naïve mESCs delays abscission through non-canonical signalling maintaining high Aurora B levels and stabilising microtubules. The influence of Wnt signalling on abscission duration is a single-cell property, that depends on cell state. Early during exit from naïve pluripotency, Wnt signalling favours delayed abscission. As cells progress toward differentiation, however, similar signalling inputs activate alternative regulatory pathways, causing fast abscission. Therefore, the role of Wnt signalling in abscission appears to be dynamically reprogrammed as naïve pluripotent cells exit naïve pluripotency, showing that non-canonical Wnt, similarly to canonical Wnt, is context dependent.

## Results

### GSK-3β activity delays abscission in naïve mESCs

We use mouse embryonic stem cells as a model to study the effect of Wnt signalling on abscission duration. By comparing naïve cells (grown in 2i-Lif media or 1i-Lif media with no GSK-3β inhibitor) and exiting cells (grown in N2B27 media), we can compare the same cell line undergoing either slow or fast abscission (Figure 1a).

**Figure 1.**
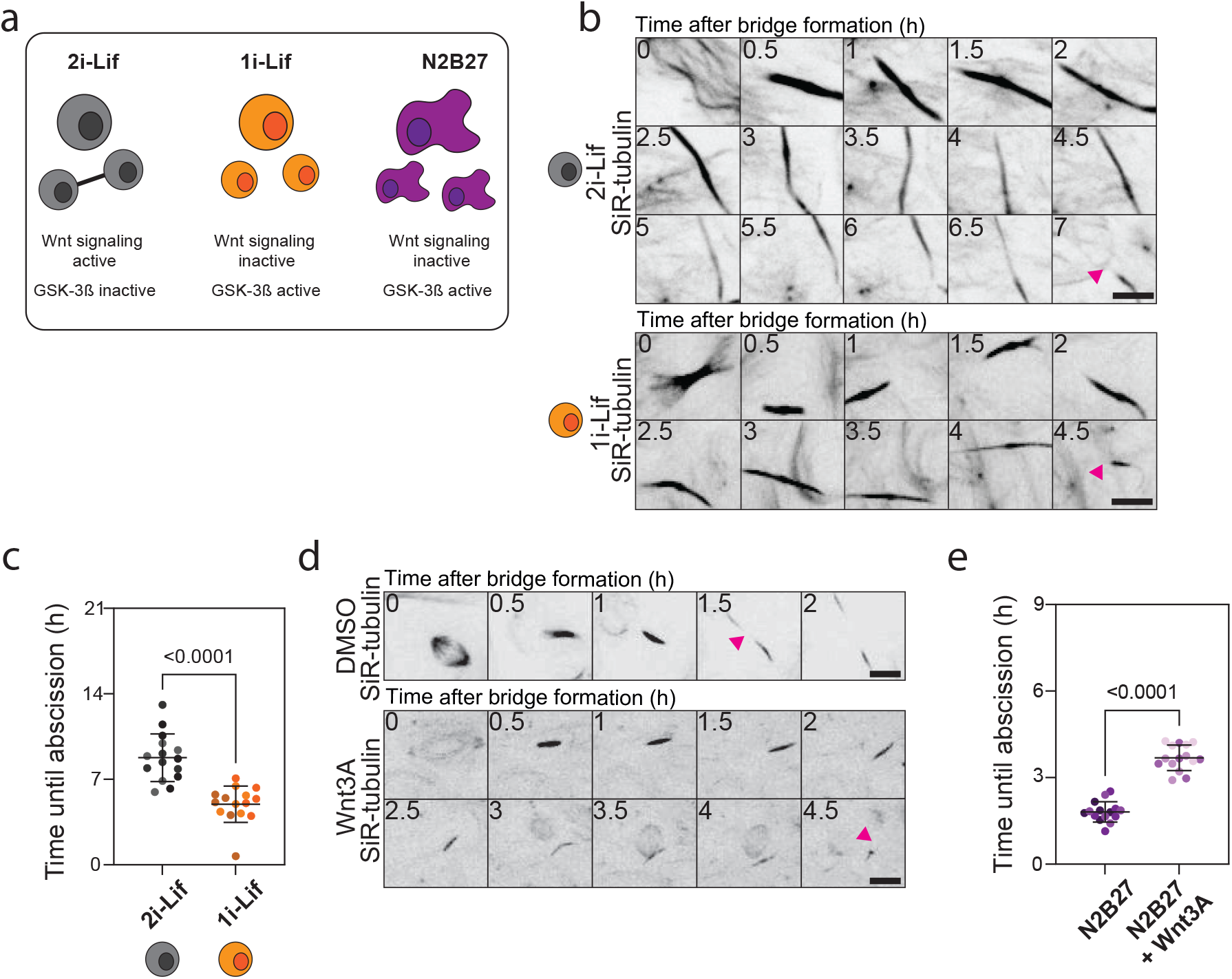
Wnt signalling delays abscission in mouse embryonic stem cells. a) Cartoon showing different conditions of growing mouse embryo9nic stem cells. Cells were grown either in 2i-Lif (grey), “1i-Lif” (2i-Lif without Chiron) (orange) or N2B27 (purple) media. b) Live-cell imaging of mESCs in 2i-LIF (top) and 1i-LIF (bottom) incubated overnight with 20 nM SiR-tubulin. A Z-projection over the height of the whole cell is shown. Tubulin is shown in black. The pink arrows indicate the cut sites. One frame is shown every 0.5h. Scale bars: 10 μm. c) Quantification of the duration of abscission from bridge formation until microtubule severing in mESCs in 2i-Lif and without Chiron. The mean and standard deviation are shown. N=3 replicates. d) Live-cell imaging of mESCs in N2B27 with DMSO (top) or Wnt3A ligand (bottom) incubated overnight with 20 nM SiR-tubulin. A Z-projection over the height of the whole cell is shown. Tubulin is shown in black. The pink arrows indicate the cut sites. One frame is shown every 0.5h. Scale bars: 10 μm. e) Quantification of the duration of abscission from bridge formation until microtubule severing in mESCs in N2B27 with (dark purple) or without (light purple) the addition Wnt3A ligand. The mean and standard deviation are shown. N=3 replicates.

mESCs are maintained in their naïve pluripotent state by inhibition of GSK-3β (which phosphorylates β-Catenin and other substrates and targets them for destruction) through the inhibitor CHIR-99021. To test if Wnt signalling plays a role in regulating abscission dynamics, we decreased Wnt signalling activity in mESCs by growing naïve ESCs without CHIR-99021 (“1i-Lif”). By using microtubule staining as a proxy for the bridges, we find that removal of CHIR-99021 leads to a strong decrease in bridge number (Supplementary Figure 1a,b), suggesting accelerated abscission. Importantly, removal of CHIR-99021 does not impair the pluripotency state of the cells nor their ability to proliferate (Supplementary Figure 1c-e). To confirm that Wnt signalling delays abscission, we imaged naïve mESCs incubated with the live microtubule probe SiR-tubulin with or without CHIR-99021 from anaphase onset to the moment of microtubule severing. We use the rupture of the microtubule bundle as a proxy for abscission, as membrane rupture promptly follows the rupture of the microtubule bundle in mESC (Kodba et al., 2024). Cells in 2i-Lif cut the cytoplasmic bridge in 10h while cells grown in 1i-Lif are almost twice as fast (Figure 1b,c, Movie 1). These results suggest that the activity of GSK-3β accelerates abscission in naïve pluripotent mESCs.

### Activation of Wnt signalling delays abscission

To test that activation of Wnt leads to slower abscission, we treated mESCs with the Dickkopf-1 (Dkk-1) inhibitor WAY-2626211 (“WAY”). Upon WAY-2626211 treatment in 1i-LIF and N2B27, the number of bridges increases (Supplementary figure 1f,g), suggesting that abscission is slower. On the other hand, the number of bridges does not change in 2i-Lif media upon Dkk-1 inhibition (Supplementary Figure 1f,g). Slowing down of abscission after Dkk-1 inhibition is also seen while live imaging mESCs exiting naïve pluripotency with WAY-262611 (Supplementary Figure 1h,i). This suggests that activation of Wnt signalling in exiting cells leads to slower abscission.

To directly activate Wnt signalling, we then treated mESCs activated Wnt signalling during exit from naïve pluripotency with the Wnt3A ligand (the Wnt ligand crucial in early mammalian development) (Barrow et al., 2003; Liu et al., 1999), which binds to Fzd receptor and activates Wnt signalling. Wnt3A addition leads to an increase in bridge number in mESC in 1i-Lif and N2B27, suggesting that abscission is delayed (Supplementary figure 2a,b). To confirm this, we imaged mESCs grown in N2B27 incubated with SiR-tubulin from anaphase onset to the moment of microtubule severing with or without Wnt3a ligand. Wnt signalling activation leads to slower abscission (Figure 1d,e, Movie 2). The cells in N2B27 exited pluripotency despite Wnt3A ligand addition (Supplementary Figure 2c-e). Altogether, these results show that activation of Wnt signalling in mESCs causes slower abscission.

### The consequences of GSK-3β inactivation in abscission of mESCs are cell state dependent

To check if the effect of GSK-3β activity on abscission depends on cell state, we treated mESCs with the GSK-3β inhibitor CHIR-99021 at different times during exit from pluripotency: at the time they started exiting naïve pluripotency or after 48 hours of exiting naïve pluripotency where cells have reached a primed state (Kalkan et al., 2017; Kinoshita et al., 2021; Kinoshita and Smith, 2018b). When mESCs exit from pluripotency in the presence of the GSK-3β inhibitor, abscission is slower than in cells in N2B27 (Figure 2a,b). Surprisingly, when CHIR-99021 is added to mESCs after 48 hours of exit from naïve pluripotency, abscission is faster than in control cells exiting naïve pluripotency (Figure 2c,d). These results suggest that the effect of GSK-3β activity on abscission duration in mESCs depends on cell state.

**Figure 2.**
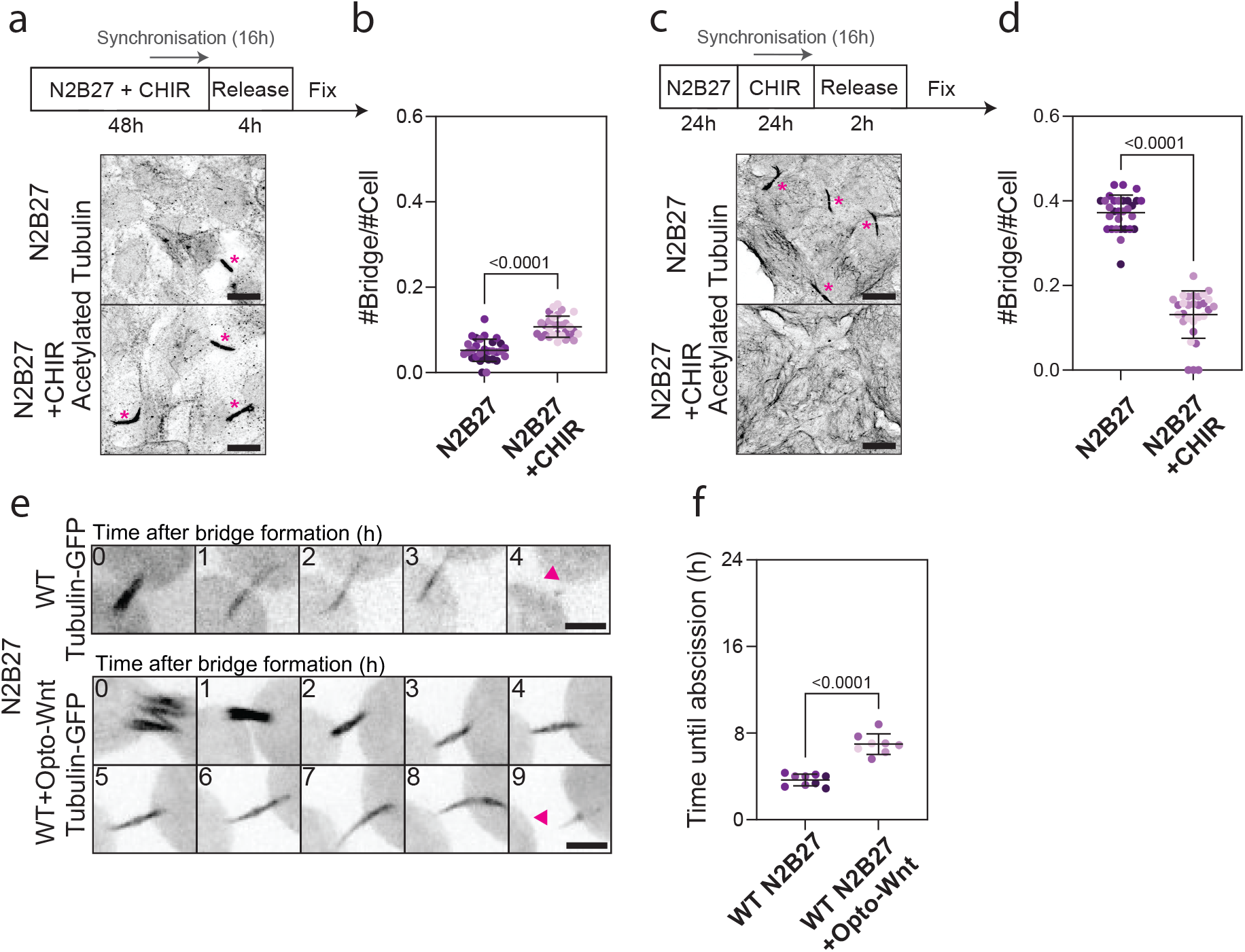
GSK-3β influence on abscission duration in mESCs depends on cell state. a) Top panel: experimental set-up of the experiment. Bottom panel: immunofluorescence showing the number of bridges in mESCs cultured in N2B27 (top) and mESCs grown in N2B27 with addition of CHIR-99021 at the beginning of exit from pluripotency (bottom). Cells were synchronized and fixed after approximately 3h spent in cytokinesis. A Z-projection over the height of the whole cell is shown. The bridges are shown with the staining of tyrosinated tubulin. Bridges are highlighted with pink asterisks. Scale bars: 10 μm. b) Quantification of the number of bridges per cell in N2B27 media (dark purple) and N2B27 with CHIR-99021 added at start of exit from pluripotency (light purple). The mean and standard deviation are shown. N=3 replicates. c) Top panel: experimental set-up of the experiment. Bottom panel: immunofluo-rescence showing the number of bridges in N2B27 grown mESCs (top) and mESCs grown in N2B27 with addition of CHIR-99021 48h after the start of exit from pluripotency (bottom). Cells were synchronized and fixed after approximately 1h spent in cytokinesis. A Z-projection over the height of the whole cell is shown. The bridges are shown with the staining of tyrosinated tubulin. Bridges are highlighted with pink asterisks. Scale bars: 10 μm. d) Quantification of the number of bridges per cell in N2B27 (dark purple) and N2B27 with CHIR-99021 added 48h after the start of exit from pluripotency (light purple). The mean and standard deviation are shown. N=3 replicates. e) Live-cell imaging of mESCs transfected with only Tubulin-GFP or Tubulin-GFP and Opto-Wnt (LRP6-Cry-mCherry). A Z-projection over the height of the whole cell is shown. Tubulin is shown in black. The pink arrows indicate the cut sites. One frame is shown every 1h. Scale bars: 10 μm. f) Quantification of the duration of abscission from bridge formation until microtubule severing in mESCs in N2B27 untransfected (WT dark purple) or expressing Opto-Wnt (LRP6-Cry-mCherry) (light purple). The mean and standard deviation are shown. N=3 replicates.

### Optogenetic activation of Wnt signalling delays abscission in exiting mESCs

Since global activation of Wnt signalling in exiting mESCs decreases abscission duration, we wanted to test whether Wnt signalling at the single cell level impacts abscission duration in mESCs. To test that, we imaged mESCs expressing Opto-Wnt (Bugaj et al., 2013; Repina et al., 2023) during exit from naïve pluripotency. Opto-Wnt contains the intracellular part of the receptor LRP6 coupled to the Cry2 protein, which undergoes oligomerization when activated with blue light, leading to clustering of the LRP6 receptor and activation of Wnt signalling. Of note, here, both control cells and cells expressing Opto-Wnt are exposed to blue light. We confirmed that Wnt signalling is activated by imaging 7xTCF-GFP (which uses 7 tandem repeats of the TCF/LEF binding site to drive expression of GFP as a reporter gene). We see that upon blue light illumination, 7xTCF-GFP is expressed in a time-scale compatible with abscission regulation (Supplementary Figure 2f). In illuminated cells expressing Opto-Wnt, abscission is slower compared to the illuminated control cells (Figure 2e,f, Movie 3). In conclusion, local activation of Wnt signalling at the single cell level is sufficient to delay abscission in mESCs.

### β-Catenin does not influence abscission duration

We then investigated how Wnt signalling controls abscission duration. To test whether canonical signalling is involved, we imaged β-Catenin KO mESCs (Wray et al., 2011) and compared abscission duration to WT mESCs. To verify the β-Catenin KO line, we first stained the cells with total and active β-catenin antibodies (Supplementary Figure 3a-c). These data show that β-Catenin is indeed absent from the β-Catenin KO line. The β-Catenin KO line retains full colony-forming potency and normal exit from pluripotency capacity (Supplementary Figure 3d-f), suggesting unaltered pluripotency, in line with studies with a full-length β-Catenin KO (Aulicino et al., 2020). To test the role of β-Catenin on abscission duration, we performed live imaging of WT and β-Catenin KO cells incubated with SiR-tubulin. Abscission in β-Catenin KO cells lasts around 6 hours, comparable to the duration of abscission in WT mESCs (Figure 3a,b, Movie 4). The number of bridges in fixed samples is also similar between β-Catenin KO and WT mESCs (Figure 3c,d). Altogether, these data show that β-Catenin does not influence abscission duration in mESCs.

**Figure 3.**
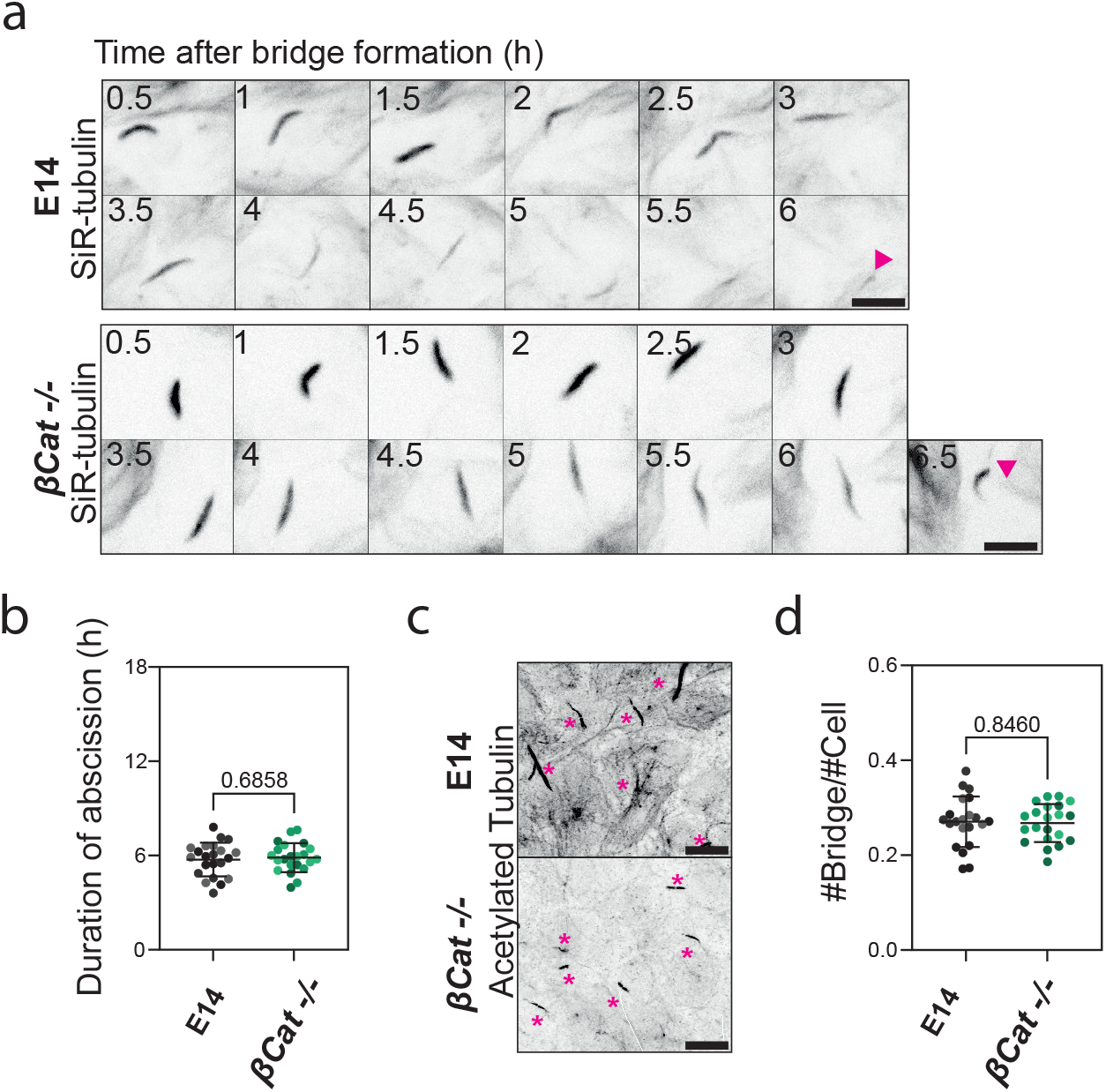
Canonical Wnt signalling does not influence abscission dynamics in mouse embryonic stem cells. a) Live-cell imaging of WT (top) and β-Catenin KO mESCs (bottom) incubated overnight with 20 nM SiR-tubulin. A Z-projection over the height of the whole cell is shown. Tubulin is shown in black. The pink arrows indicate the cut sites. One frame is shown every 0.5h. Scale bars: 10 μm. b) Quantification of the duration of abscission from bridge formation until microtubule severing in mESCs in WT (grey) and β-Catenin KO (green) mESCs. The mean and standard deviation are shown. N=3 replicates. c) Immunofluorescence showing the number of bridges in WT (top) and β-Catenin KO mESCs (bottom). A Z-projection over the height of the whole sample is shown. The bridges are shown with the staining of tyrosinated tubulin. Bridges are highlighted with pink asterisks. Scale bars: 10 μm. d) Quantification of the number of bridges per cell in WT (grey) and β-Catenin KO mESCs (green). The mean and standard deviation are shown. N=3 replicates.

### GSK-3β controls microtubule stability at the bridge of mESC

Since non-canonical signalling is not involved in abscission regulation, we tested non canonical function of GSK-3β. GSK-3β can phosphorylate microtubule-binding proteins and influence microtubule stability, a key player in abscission dynamics (Kodba et al., 2024). Therefore, we tested if GSK-3β activity influences microtubule stability at the bridge. First, we mapped the localization of GSK-3β during bridge maturation. GSK-3β localizes at the midbody throughout bridge maturation (Figure 4a, Movie 5), at similar endogenous levels whether GSK-3β is active or not (Figure 4b,c). The microtubules at the bridge in mESCs in 1i-Lif media are shorter than microtubules in the bridge in 2i-Lif media at the same timepoint during bridge maturation (Supplementary Figure 4a,b). We then assessed microtubule stability using post-transcriptional modifications of tubulin as a proxy; indeed, stable microtubules tend to be acetylated (Janke and Montagnac, 2017). Our data show that mESCs with active GSK-3β contain less acetylated microtubules at the bridge compared to cells with GSK-3β inhibitors at the same timepoint during cytokinesis (Figure 4d,e). Of note, we normalize the amount of acetylated tubulin by bridge size to take into consideration that the bridges are shorter in 1i-Lif compared to 2i-Lif media. However, the amount of total tubulin at the bridge (also normalized by bridge size) is the same whether GSK-3β is active or inactive (Figure 4f,g). Altogether, our data show that GSK-3β activity reduces the size of the microtubule and their acetylation at the bridge, suggesting that GSK-3β activity decreases microtubule stability at the bridge.

**Figure 4.**
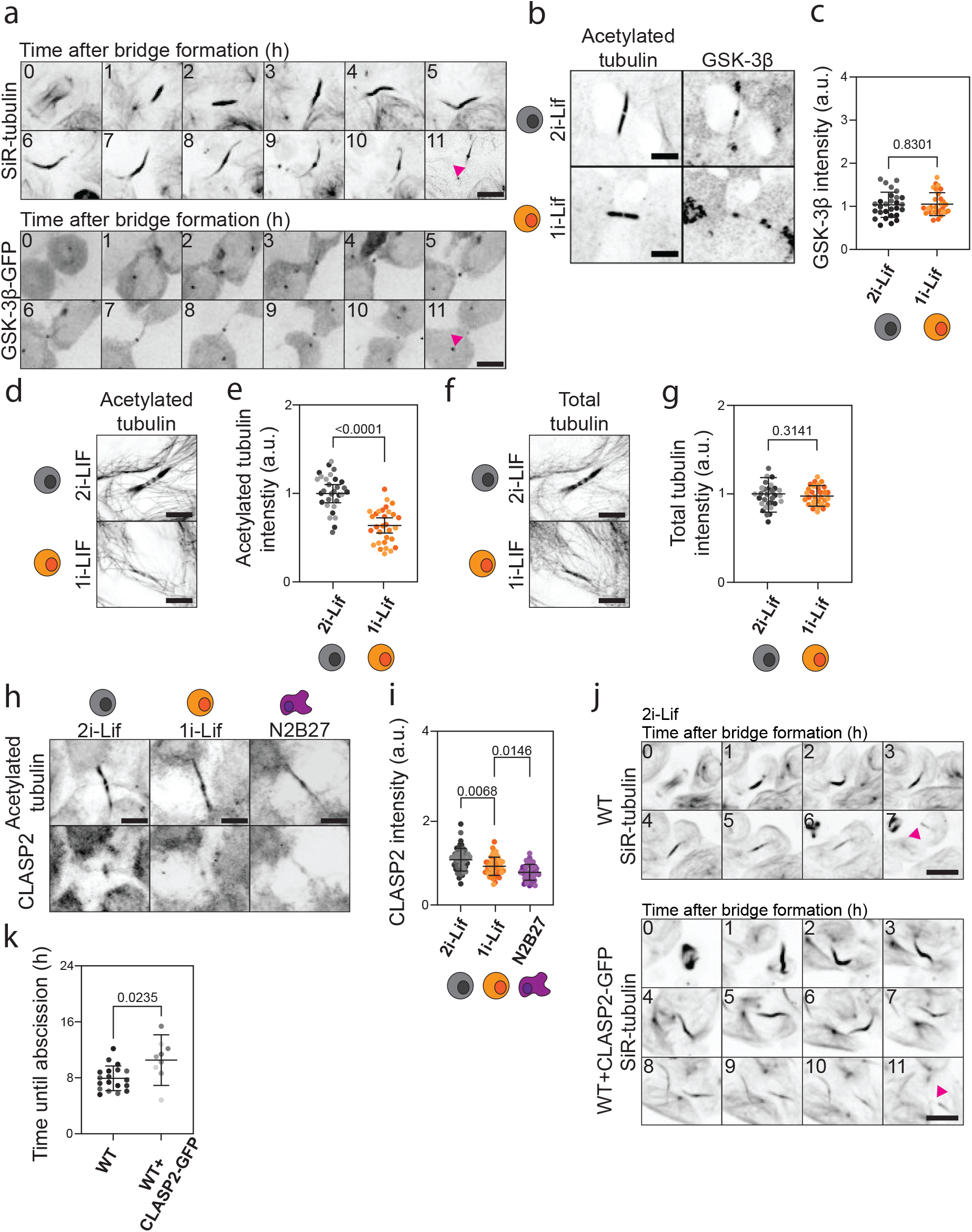
Wnt signalling affects microtubules and their binding proteins. a) Live-cell imaging of mESCs in 2i-Lif media incubated overnight with 20 nM SiR-tubulin (top) and transfected with GSK-3β-GFP (bottom). A Z-projection over the height of the whole cell is shown. The pink arrows indicate the cut sites. One frame is shown every 1h. Scale bars: 5 μm. b) Immunofluorescence showing the localization of acetylated tubulin (left) and GSK-3β (right) at the bridge in 2i-Lif (top) and 1i-Lif (bottom) media. Scale bar: 5 μm. c) Quantification of the intensity of GSK-3β in mESCs grown in 2i-Lif (grey) and 1i-Lif (orange). The mean and standard deviation are shown. N=3 replicates. d) Immunofluorescence showing the localization of acetylated tubulin at the bridge in 2i-Lif (top) and 1i-Lif (bottom). Scale bar: 5 μm. e) Quantification of the intensity of acetylated tubulin in mESCs in 2i-Lif (grey) and 1i-Lif (orange) media. The mean and standard deviation are shown. N=3 replicates. f) Immunofluorescence showing the localization of total tubulin at the bridge in 2i-Lif (top) and 1i-Lif (bottom) media. Scale bar: 5 μm. g) Quantification of the intensity of total tubulin in mESCs in 2i-Lif (grey) and 1i-Lif (orange). The mean and standard deviation are shown. N=3 replicates. h) Immunofluorescence showing the localization of CLASP2 at the bridge in 2i-Lif, 1i-Lif and N2B27 (bottom row, from left to right, respectively). A Z-projection over the height of the whole sample is shown. The bridges are shown with the staining of acetylated tubulin. Scale bar: 5 μm. i) Quantification of CLASP2 intensity in mESCs in 2i-Lif (grey), 1i-Lif (orange) and N2B27 (purple). The mean and standard deviation are shown. N=3 replicates. j) Live-cell imaging of WT (top) and WT+CLASP2-GFP (bottom) incubated overnight with 20 nM SiR-tubulin in 2i-Lif. A Z-projection over the height of the whole cell is shown. Tubulin is shown in black. The pink arrows indicate the cut sites. One frame is shown every 1h. Scale bars: 10 μm. k) Quantification of the duration of abscission from bridge formation until microtubule severing in mESCs in WT (dark grey) and WT+CLASP2-GFP (light grey) mESCs. The mean and standard deviation are shown. N=3 replicates.

### CLASP2 delays abscission in mESC

GSK-3β kinase could directly phosphorylate microtubules associated proteins (MAPs) at the bridge, which in turn could locally regulate microtubule stability. CLIP-associating proteins (CLASPs) 1 and 2 are GSK-3β targets (Kumar et al., 2009). First, we tested whether CLASP1 and CLASP2 localize at the bridge of mESC. CLASP1 and CLASP2 localize at the intracellular bridge of mESC, with CLASP1 localizing across the whole bridge (Supplementary Figure 4c,d) and CLASP2 specifically localizing at the midbody (Figure 4h,i). A time-course analysis revealed that while CLASP1 is along the bridge during bridge maturation (Supplementary Figure 4e,f), CLASP2 distributes along the length of the bridge during bridge formation but is progressively restricted to the midbody during bridge maturation (Supplementary Figure 4g,h). The amount of both CLASP1 and CLASP2 decreases over time during bridge maturation (Supplementary Figure 4e-h). Importantly, the amount of both CLASP1 and CLASP2 depends on the activity of GSK-3β, with higher CLASP1/2 at the bridge when GSK-3β is inactive (2i-Lif condition, Figure 4h,i and Supplementary Figure 4c,d).

Because the localization of CLASP2 is similar the localization of GSK-3β at the bridge, and GSK-3β activity decreases the amount of acetylated microtubules, the amount of CLASP2, and accelerates abscission, we hypothesized that GSK-3β could regulate the duration of abscission by modulating microtubule dynamics at the midbody through CLASP2. To test this, we overexpressed CLASP2 in mESCs with or without the inhibitor of GSK-3β. In 1i-Lif, where GSK-3β is active, the overexpression of CLASP2 has no effect on abscission dynamics (Supplementary Figure 4i,j). However, in 2i-Lif, where GSK-3β is inactive, the overexpression of CLASP2 delays abscission by 27% (Figure 4j,k, Movie 6). In conclusion, CLASP2 delays abscission in absence of GSK-3β activity.

### Preventing protein degradation leads to delayed abscission

GSK-3β activity controls abscission dynamics through effects on microtubules, possibly through CLASP1/2, independently of β-Catenin; however, this does not mean that other targets of GSK-3β are not also involved. In particular, Wnt-dependent stabilization of proteins (Wnt/STOP) which is active during mitosis (Acebron et al., 2014) could be involved in regulation of abscission. To test whether protein degradation regulates abscission dynamics, we synchronised mESCs after 48h of exit from naïve pluripotency and treated them with a proteasome inhibitor (MG132) after bridge formation. To confirm that MG132 decreases protein degradation, we stained mESCs with an ubiquitin antibody that specifically recognizes ubiquitin-remnant motif (K-Epsilon-GG) (Draczkowski et al., 2025; Tsuchiya et al., 2017) after MG132 treatment and confirmed that there is indeed more ubiquitin at the bridges after MG132 treatment (Supplementary Figure 5a,b). Furthermore, preventing protein degradation significantly delays abscission since the number of bridges increases upon MG132 treatment (Figure 5a,b). These data show that protein degradation leads to faster abscission.

**Figure 5.**
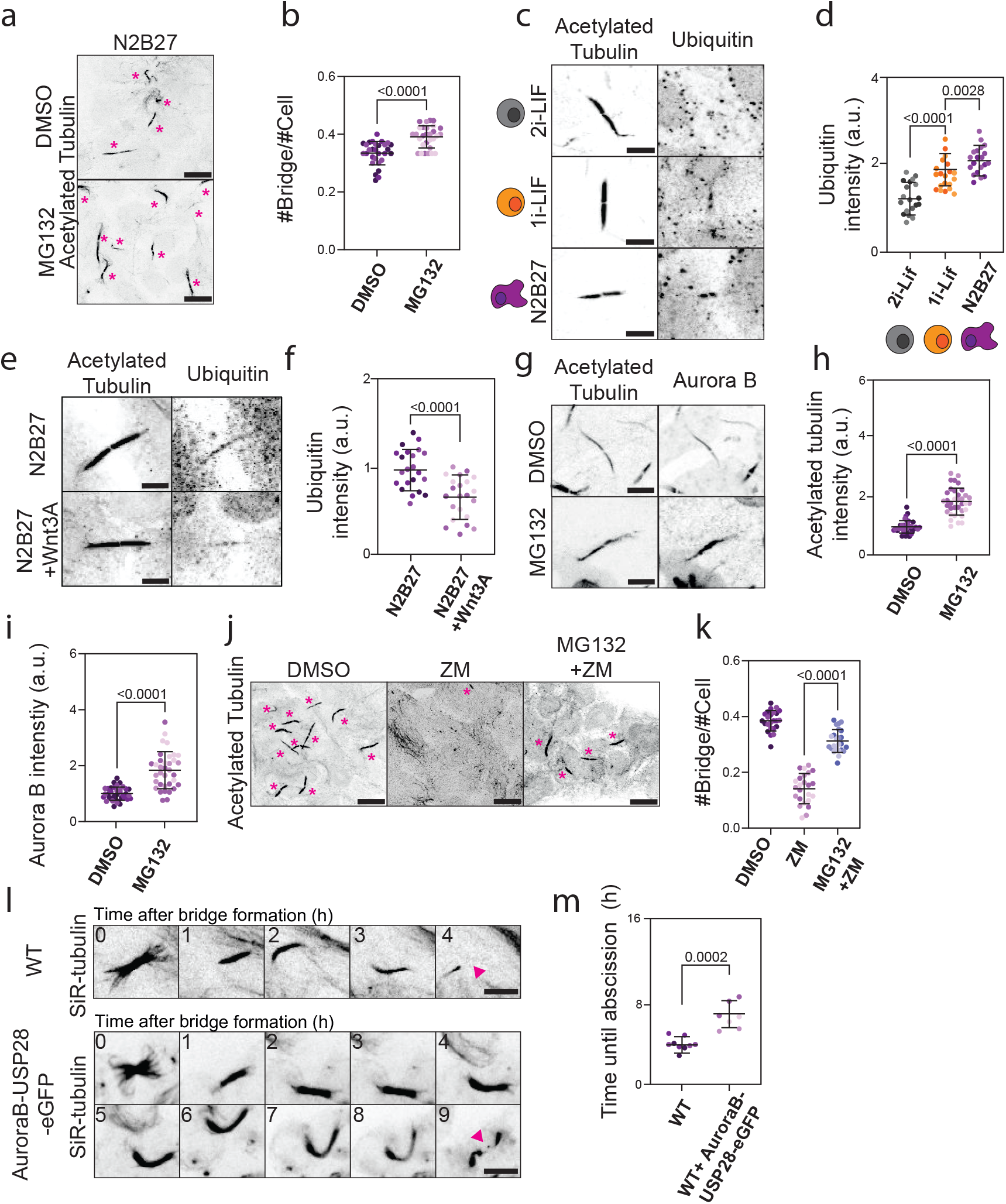
Protein degradation leads to faster abscission. a) Top: experimental set-up of proteosome inhibition experiments. Bottom: immunofluorescence showing the number of bridges in control, DMSO treated (top) and MG132 treated (bottom) cells in N2B27. A Z-projection over the height of the whole sample is shown. The bridges are shown with the staining of acetylated tubulin. Bridges are highlighted with pink asterisks. Scale bars: 10 μm. b) Quantification of the number of bridges per cell in control, DMSO treated (grey) and MG132 treated (orange) cells in N2B27. The mean and standard deviation are shown. N=3 replicates. c) Immunofluorescence showing the localization of ubiquitin at the bridge in 2i-Lif, 1i-Lif and N2B27. A Z-projection over the height of the whole sample is shown. The bridges are shown with the staining of acetylated tubulin. Scale bar: 10 μm. d) Quantification of ubiquitin intensity in mESCs in 2i-Lif (grey), 1i-Lif (orange) and N2B27 (purple). The mean and standard deviation are shown. N=3 replicates. e) Immunofluorescence showing the localization of ubiquitin at the bridge in N2B27(top) or with addition of Wnt3A ligand (bottom). A Z-projection over the height of the whole sample is shown. The bridges are shown with the staining of acetylated tubulin. Scale bar: 10 μm. f) Quantification of ubiquitin intensity in mESCs grown in N2B27 (dark purple) or with the addition of Wnt3A ligand (light purple). The mean and standard deviation are shown. N=3 replicates. g) Immunofluorescence showing the localization of Aurora B and acetylated tubulin at the bridge in mESCs in N2B27 treated with DMSO (top) or MG132 (bottom). A Z-projection over the height of the whole sample is shown. Scale bar: 10 μm. h) Quantification of acetylated tubulin intensity in mESCs in N2B27 treated with DMSO (dark purple), or MG132 (light purple). The mean and standard deviation are shown. N=3 replicates. i) Quantification of Aurora B intensity in mESCs in N2B27 treated with DMSO (dark purple), or MG132 (light purple). The mean and standard deviation are shown. N=3 replicates. j) Immunofluorescence showing the number of bridges in DMSO (left), ZM (middle) and MG132+ZM (right) treated mESCs in N2B27. A Z-projection over the height of the whole sample is shown. The bridges are shown with the staining of acetylated tubulin. Bridges are highlighted with pink asterisks. Scale bars: 10 μm. k) Quantification of the number of bridges per cell in DMSO (dark purple), ZM (light purple) and MG132+ZM (blue) treated cells in N2B27. The mean and standard deviation are shown. N=3 replicates. l) Live-cell imaging of WT (top) and WT+AuroraB-USP28-eGFP (bottom) incubated overnight with 20 nM SiR-tubulin in N2B27. A Z-projection over the height of the whole cell is shown. Tubulin is shown in black. The pink arrows indicate the cut sites. One frame is shown every 0.5h. Scale bars: 10 μm. m) Quantification of the duration of abscission from bridge formation until microtubule severing in mESCs in WT (dark purple) and WT+AuroraB-USP28-eGFP (light purple) grown in N2B27. The mean and standard deviation are shown. N=3 replicates.

### Active Wnt signalling decreases ubiquitinylation at the bridge

To test whether there are intrinsic differences in protein degradation when Wnt signalling is active, we examined our ubiquitin staining in conditions with active or inactive GSK-3β. Our data show that in all conditions, ubiquitin-remnant motif (K-Epsilon-GG) is present at the bridge (Figure 5c,d, Supplementary Figure 5a,b). When GSK-3β is active (1i-Lif and N2B27), there is more ubiquitin-remnant motif at the bridge (Figure 5c,d, Supplementary Figure 5a,b). This suggests that there is intrinsically higher activity of protein degradation through ubiquitin at the bridge when Wnt signalling is inactive. To confirm this, we treated cells with Wnt3A and stained for ubiquitin-remnant motif. When the GSK-3β inhibitor is not present (N2B27), the addition of Wnt3A leads to less ubiquitin-remnant motif present at the bridge (Figure 5e,f). Altogether, our data suggest that active Wnt signalling leads to decreased ubiquitinylation at the bridge.

### Protein degradation leads to less Aurora B and acetylated microtubules at the bridge

Microtubule stability and Aurora B activity at the bridge are major drivers of slow abscission (Kodba et al., 2024). Therefore, we wanted to test the effects of protein degradation on microtubule acetylation, which we use as a proxy for microtubule stability, and Aurora B levels. Upon addition of MG132 in exiting mESCs, we see an increased amount of acetylated microtubules at the bridge (Figure 5g,h) as well an increase in the amount of Aurora B at the bridge (Figure 5g,i). The amount of total tubulin at the bridge remains the same upon MG132 treatment as in control cells (Supplementary Figure 5c,d). The intensity of tubulin and Aurora B is normalised by the bridge size. These data indicate that the activity of the proteosome controls the amount of Aurora B kinase and acetylated microtubules at the bridge. Finally, we tested whether the inhibition of GSK-3β could recapitulate the effects of proteasome inhibition. Indeed, in cells where the activity of GSK-3β was inhibited (2i-Lif), there was more Aurora B and P-Aurora B at the bridge (Supplementary Figure 5e,f). Altogether, these data suggest that Aurora B levels and microtubule acetylation are controlled by protein degradation under the dependence of GSK-3β activity.

### Preventing Aurora B degradation leads to slow abscission

Our previous data show that the activity of Aurora B kinase at the bridge promotes slow abscission and that the amount of Aurora B at the bridge is increased upon proteosome inhibition. Thus, we tested whether reducing Aurora B activity would rescue proteasomal inhibition. Indeed, treating mESCs with both Aurora B inhibitor and MG132 largely rescues slow abscission (Figure 5j,k), suggesting that abscission duration is controlled by the degradation of Aurora B at the bridge. To test this, we designed a construct where Aurora B kinase was fused to USP28 deubiquitinylase to constantly prevent its ubiquitination and subsequent degradation (Stringer and Piper, 2011) (see Methods). We first checked that Aurora B-USP28-eGFP persists at the bridge. We observe that Aurora B-USP28-eGFP is present at the bridge until abscission in all cells (Supplementary Figure 5h). On the other hand, in cells transfected with INCENP-GFP, the binding partner of Aurora B, INCENP disappears from the bridge 60 ± 20 min before abscission (15 cells from 3 different experiments, Supplementary Figure 5h). These data suggest that the Aurora B-USP28-eGFP remains at the bridge longer than the endogenous Aurora B. Furthermore, in the absence of Wnt signalling, cells expressing Aurora B-USP28-eGFP have slower abscission than control cells (Figure 5l,m, Movie 7). In conclusion, preventing Aurora B ubiquitinylation leads to its maintenance at the bridge and slow abscission in mESCs.

## Discussion

In this paper, we investigate how Wnt signalling regulates abscission duration in mESCs. We show that active Wnt signalling delays abscission in mESCs. Our data, together with reports that Wnt signalling plays a role in cytokinesis (Darmasaputra et al., 2024; Fumoto et al., 2012), pinpoint Wnt as an important regulator of the last stages of cell division, notably through its kinase GSK-3β (Figure 6, top). Beside the direct manipulation of GSK-3β activity, we also use the Wnt ligand Wnt3A and Opto-Wnt. Both of these treatments recapitulate the slow abscission, probably by also deactivating GSK-3β as a downstream target of Wnt signalling activation. While this showed the global effect of Wnt signalling on abscission dynamics, we wanted to also test whether locally activating Wnt signalling in mESCs grown in N2B27 leads to slower abscission. By using precise optogenetic Wnt activation, we show that local activation of Wnt signalling also results in slow abscission and that abscission dynamics is a property at the single-cell level.

**Figure 6.**
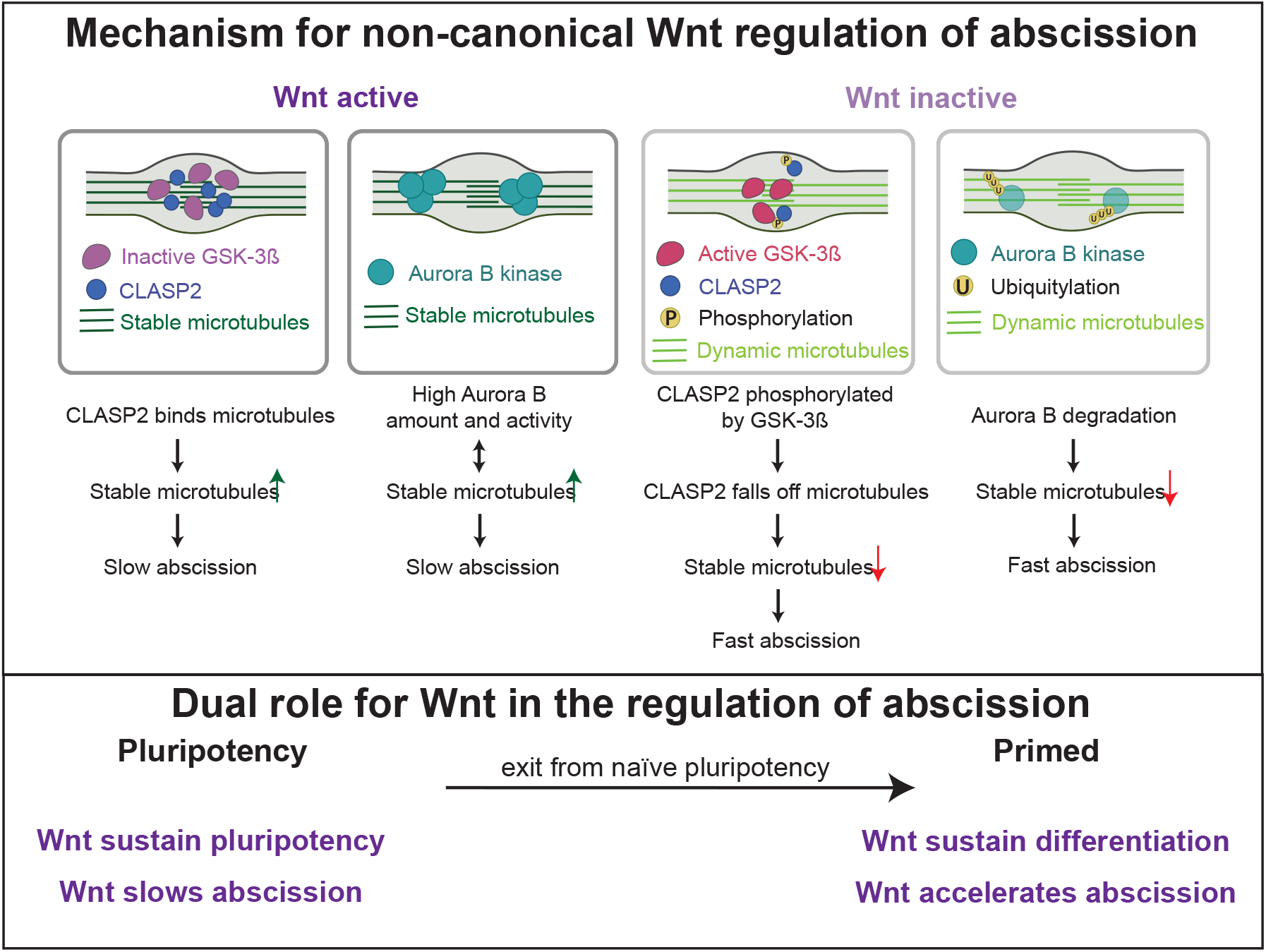
Proposed model for regulation of abscission dynamics in mouse embryonic stem cells. When Wnt signalling is active, there is more CLASP2 at the bridge. Since CLASP2 acts as microtubule stabilizer, we propose that CLASP2 binding to microtubules leads to more stable microtubules at the bridge. More stable microtubules at the bridge lead to slow abscission. When Wnt signalling is active, GSK-3ß is inactive and cannot phosphorylate CLASP2 at the bridge which increases its affinity for microtubules at the bridge. When Wnt is inactive, we propose that CLASP2 has lower affinity for the microtubules, leading to fewer stable microtubules at the bridge and fast abscission. In parallel, we propose that the amount of Aurora B at the bridge is regulated by GSK-3ß phosphorylation. When Wnt signalling is active, GSK-3ß is inactive and therefore cannot phosphorylate Aurora B which prevents consequent Aurora B degradation at the bridge. Conversely, active GSK-3ß would phosphorylate Aurora B which would target it for degradation, leading to faster abscission through microtubule regulation. Finally, the role of Wnt signalling on abscission is context dependent.

We show that the regulation of abscission dynamics in mESCs involves protein degradation under the dependence of GSK-3β. When GSK-3β is inactive, protein ubiquitination at the bridge is overall lower, which probably allows proteins to stay at the intercellular bridge for longer time. High ubiquitination of proteins at the bridge correlates with delayed abscission. In particular, the abundance of the abscission master regulator Aurora B at the bridge depends both on GSK-3β inhibition and proteasome degradation, suggesting that regulating the ubiquitination of Aurora B could determine the duration of abscission, consistent with prior data (Halcrow et al., 2022). Remarkably, fusing Aurora B to a deubiquitinylase slows abscission down, strengthening the hypothesis that Aurora B ubiquitinylation leads to its degradation and a faster abscission. It is tempting to hypothesize that GSK-3β directly phosphorylates Aurora B to target it for degradation, as both Aurora B and its binding partner Incenp possess the consensus motif (Ser/Thr) with a possible non-canonical priming site, common for GSK-3β substrates (Sutherland, 2011b). Further studies will be necessary to test this hypothesis and understand whether the putative regulation of Aurora B by GSK-3β is local (at the bridge) or at the cell level. It would be interesting to also test whether more abscission regulators are under control of protein degradation or GSK-3β activity. Interestingly, proteasomal activity also crosstalks with cell fate: in mESC, the deubiquitinase Psmd14 is required for maintaining pluripotency (Buckley et al., 2012), exit from naïve pluripotency coincides with an increased degradation of damaged proteins by the proteasome (Hernebring et al., 2013, 2006), while human ESC display higher proteasomal activity their differentiating counterpart (Vilchez et al., 2012). In the case of abscission, it is not yet clear whether the increase in ubiquitinylation when Wnt is not active is a direct consequence of Wnt activity, or a downstream effect of cell fate. Conversely, it will be important to test whether increased/decreased ubiquitinylation and degradation are regulated locally at the bridge, or are a global property of the cell.

We also show that GSK-3β activity influences microtubule acetylation at the bridge. This could be a direct effect of GSK-3β on microtubule regulating proteins or an indirect effect of the stabilization of Aurora B at the bridge, as we have shown previously that high Aurora B activity leads to microtubule stability (and thus acetylation) in mESC (Kodba et al., 2024). Active GSK-3β phosphorylates various microtubule interacting proteins and changes their preference for binding microtubules. This is particularly important in migrating cells where GSK-3β activity regulates microtubule architecture and polarized movement of cells by regulating microtubule crosslinking protein ACF7 (Yucel and Oro, 2011). Inhibition of GSK-3β also allows CLASP2 and APC to localize to plasma membrane and facilitate cell motility (Zaoui et al., 2010). In this paper we identify CLASP2 as an abscission regulator and a possible target of GSK-3β phosphorylation at the bridge. Further work will need to confirm that GSK-3β can indeed phosphorylate CLASP2 at the bridge, leading to its detachment from microtubules. Another possibility for Wnt signalling to influence abscission dynamics is by modulating cortex attachments at the bridge. For example, dimerization of LRP6 receptors upon Wnt ligand binding leads to local stabilization of actomyosin contractility through accumulation of Myosin1C (Eli et al., 2025). Therefore, activation of Wnt signalling could also lead to changed actomyosin contractility at the bridge. The role of actomyosin cortex in the duration of abscission has been studied in germ cells where actomyosin contractility regulators stabilize intercellular bridges and prevent abscission (Goupil et al., 2017), but nothing is known in mESCs. Finally, while CLASP1 and CLASP2 seem to have redundant functions in certain contexts (Mimori-Kiyosue et al., 2005; Pereira et al., 2006), they display distinct localisation after bridge formation, with CLASP1 along the whole length of the bridge and CLASP2 being restricted at the midbody (where the + ends probably localise) as the bridge matures. This suggests that CLASP1 and CLASP2 might be differently regulated and perform different functions during abscission.

In mESCs, Wnt signalling is maintained by the addition of CHIR-99021 which inhibits GSK-3β activity. By adding or removing CHIR-99021, we show that inhibiting GSK-3β is not necessary for pluripotency (consistent with previous reports that Wnt signaling supports pluripotency, but is not required for the maintenance of the pluripotency network (Aulicino et al., 2020)) but keeps abscission slow. Interestingly, inhibiting GSK-3β activity in mESCs exiting naïve pluripotency has different effects on abscission dependent on when CHIR-99021 is added to the media. Inhibiting GSK-3β when cells start exiting naïve pluripotency leads to slower abscission compared to cells with active GSK-3β. When we activate Wnt at the single cell level using optogenetics 16h after triggering exit, we also cause delayed abscission. On the other hand, inhibiting GSK-3β after mESCs have been exiting naïve pluripotency for 48 hours leads to faster abscission. These results suggest that abscission regulation depends on both Wnt signalling activity and cell state. Therefore, our results suggest that abscission duration is tightly coupled to cell state. Once cells have exited naïve pluripotency, Wnt signalling might no longer support the same mechanisms of regulation of abscission dynamics as in naïve pluripotent mouse embryonic stem cells which changes the duration of abscission as cells exit naïve pluripotency. Wnt signalling has a dual role during development. In naïve pluripotent mESCs, active Wnt signalling promotes pluripotency, but in epiblast ESCs it promotes differentiation (Muñoz-Descalzo et al., 2015; Sonavane and Willert, 2023). This dual role for Wnt is generally thought to be due to its permissive, rather than instructive, signalling activity of the canonical pathway; indeed, β-Catenin can bind multiple cofactors e.g., TCF1 or TCF3 which have different role. Furthermore, the chromatin landscape changes between naïve and primed state (Kinoshita and Smith, 2018b), potentially allowing binding of β-Catenin and partners to bind at different loci. Crucially, our data shows that in naïve mESCs, Wnt signalling control of abscission duration is β-Catenin independent, suggesting that the context-dependent action of Wnt signalling might not be limited to canonical Wnt signalling (Figure 6, bottom).

In conclusion, we show that active Wnt signalling delays abscission in mESCs. Active Wnt signalling and inactive GSK-3β lead to more stable microtubules at the bridge and slower abscission. We propose a dual model where Wnt signalling acts through GSK-3β inhibition to prevent Aurora B degradation and promote microtubule stability, both of which delay abscission. More broadly, our study suggest that multiple redundant mechanisms are at play to regulate abscission dynamics and identifies protein degradation as a key regulator of abscission duration in stem cells. Finally, our data suggests that non-canonical Wnt could also functions as a permissive, rather than instructive, pathway.

## Methods

### Data availability

All materials and data used in the analysis are available upon request. The data will be available on the data management platform Yoda of Utrecht University according to FAIR guidelines upon publication.

### Cell culture and transfection

#### Cell lines

Mouse embryonic stem cells E14 (a kind gift from Niels Geijsen, Hubrecht Institute) were used throughout the study. β-catenin KO cells (a kind gift from Austin Smith, University of Exeter) were used for Figure 3 and Supplementary Figure 3.

#### Cell culture

E14 mouse embryonic stem cells were cultured on 10 cm cell culture dishes (Greiner Bio-one, #664160) coated with 0.1% gelatin/PBS in N2B27+2i-LIF media with penicillin and streptomycin. Cells were cultured at a controlled density (1.5-3.0 10^4^ cells/cm^2^) and passaged every other day using Trypsin (Sigma-Aldrich, #A6964). The cells were kept in incubators at 37°C with 5% CO_2_. The cells were regularly tested for mycoplasma. Cells were plated in N2B27 after passaging to trigger exit from naïve pluripotency. For a typical experiment, cells were passaged and plated on laminin coated plates for 48h in either N2B27+2i-Lif (naïve), N2B27+1i-Lif (naïve) or N2B27 (exiting) media.

The culture medium was made in house as described in Chaigne et al., 2020. using DMEM/F-12, 1:1 mixture (Sigma-Aldrich, #D6421-6), Neurobasal medium (Life technologies #21103-049), 2.2 mM L-Glutamin (Thermofischer Scientific # 25030024), home-made N2 (see below), B27 (Life technologies #12587010), 3 µM Chiron (Merck #SML1046), 1 µM PD 0325901 (Merck #PZ0162), LIF (Merck # ESG1107), 0.1 mM β-Mercapto-ethanol, 12.5 µg/mL Insulin zinc (ThermoFischer Scientific # 12585014). The 200 X home-made N2 was made using 8.791 mg/mL Apotransferrin (Sigma-Aldrich #T1147), 1.688 mg/mL Putrescine (Sigma-Aldrich #P5780), 3 µM Sodium Selenite (Sigma-Aldrich #S5261), 2.08 µg/mL Progesterone (Sigma-Aldrich #P8783), and 8.8% BSA.

For fixed and live imaging, the cells were plated on 8-well Ibidi chambers (IBI Scientific #80807) coated with 10 mg/mL Laminin (Sigma #11243217001) overnight at 37°C.

#### Transfection

Approximately 3.10^4^ cells were seeded on a 1 cm^2^ well of an 8-well Ibidi chamber for transfection (IBI Scientific #80807). The Ibidi chamber was previously coated with 10 mg/mL Laminin (Sigma, #11243217001) overnight at 37°C.

For transfection, 1 µg of DNA was incubated in 50 µL Optimem (Thermofischer #11058021) for 5 min at room temperature and at the same time 1.2 µL of Lipofectamine™ (Thermofischer #18324012) was incubated at room temperature in 50 µL Optimem. The DNA and Lipofectamin solutions were combined after a 5 minute incubation and then incubated at room temperature for 20 min. Meanwhile, the cells were passaged as described above and seeded in the well from which Laminin has been removed and replaced with 100 µL of culture media (N2B27+2i-Lif or N2B27+1i-Lif for naïve cells or N2B27 for exiting cells). The transfection mix was added on top of the cells, and the cells were placed in the incubator overnight and typically rinsed the next morning and imaged in the afternoon.

The plasmids used in this study were: Tubulin-GFP, GSK-3ß-eGFP adapted from Addgene plasmid #49491 by adding a GFP at the C-terminus, Aurora B-USP28-eGFP, mCherry-Cry2-LRP6c (Repina et al., 2023), 7xTCF-GFP (Repina et al., 2023).

### Drug treatment

#### Synchronization

When indicated, cells were synchronized using 10μM RO-3306 (Selleckchem/BioConnect # S7747) for 15h and released for 90 min before further treatment. For the time course of bridge maturation, cells were fixed 2h after washing out the RO-3306 which corresponds to bridge formation (0h), or 2h or 4h later.

#### Inhibition of Dkk-1 with WAY262611

When indicated, cells were treated with 2μM WAY262611 (Selleckchem/BioConnect # S9828) or an equivalent amount of DMSO for 24 hours, then fixed or live imaged.

#### Inhibition of proteasome with MG132

When indicated, cells were treated with 5μM MG132 (Selleckchem/BioConnect # S2619) or an equivalent amount of DMSO for 90 minutes, then fixed.

#### Inhibition of Aurora B kinase activity with ZM447439

When indicated, cells were treated with 2μM ZM (Selleckchem/BioConnect # S1103) or an equivalent amount of DMSO for 90 minutes, then fixed.

#### Activation of Wnt pathway using Wnt3A

When indicated, cells were treated with 100ng/mL Wnt3A (Merck Life Science/Sigma Aldrich # GF154) or an equivalent amount of DMSO.

### Immunofluorescence

The primary antibodies used were alpha Tubulin Monoclonal Antibody (ThermoFischer Scientific #62204), P-Aurora B (ThermoFischer Scientific # PA5-105026), Aurora B AIM-1 (BD Biosciences # 611082), Acetylated Tubulin (Merck # T7451 and Cell Signalling Technology # D20G3), Tyrosinated tubulin (ThermoFischer Scientific #MA1-80017), GSK-3β (Cell Signalling Technology #9315), β-catenin (Cell Signalling Technology #9562), active β-catenin (Cell Signalling Technology #8814), K48-ubiquitin (ThermoFischer # MA5-35382).

The immunofluorescence was done as described (Kodba and Chaigne, 2023b). The cells were fixed with 4% paraformaldehyde for 10 minutes and then rinsed 3 times with PBS. Cells were then permeabilized with 0.1% Triton in PBS for 10 minutes and rinsed 3 times in PBS. Afterwards, the cells were blocked in 3% BSA in PBS for 15 minutes. Primary antibodies were added at 1:200 in the blocking solution and incubated for 2h at room temperature. Then, cells were rinsed 3 times in PBS and again blocked for 15 minutes with the blocking solution and incubated with the secondary antibodies at 1:500 for 1h at room temperature. Cells were then rinsed 3 times with PBS, incubated for 10 minutes with Hoechst, rinsed 3 times with PBS, and kept in PBS until imaging.

For imaging of fixed samples, a Zeiss LSM-700 confocal setup was used. It consists of an AxioObserver Z1 microscope with a Plan-Apochromat 520 63×/0.8 oil objective. The set-up was controlled using ZEN.

### Live imaging

For live imaging of microtubules, cells were incubated overnight with 20 nM SiR-tubulin (Spirochrome #SC002).

Live imaging was performed using a 60× (Plan Apo VC, NA 1.4; Nikon) oil-immersion objective on a Spinning Disc (Yokogawa CSU-X1-A1) Nikon Eclipse Ti microscope with Perfect Focus System equipped with a sample incubator (Tokai-Hit) and an Evolve 512 EMCCD camera (Photometrics), controlled with MetaMorph 7.7 software (Molecular Devices). Cobolt Calypso 491 nm and Cobolt 647 nm lasers were used for excitation. Images were acquired every 5 min.

### Optogenetic stimulation

Cells were seeded on a 1 cm^2^ well of an 8-well Ibidi chamber and placed on Spinning Disc (Yokogawa CSU-X1-A1) Nikon Eclipse Ti microscope around 16 hours after seeding the cells. Optogenetic stimulation was achieved with blue light emitted by 470 nm laser illuminating mESCs every 30 minutes with a pulse of blue light for the duration of the experiment (12-24 h).

### Colony-forming assay

Colony-forming assays were done to test the dynamics of exit from naïve pluripotency. Cells were plated at low density (3×10^4^ cells per well of a 24-well plate) on plates previously coated with 0.1% gelatin in N2B27 for 20 hours. After 20h, cells were resuspended, counted, and plated at clonal density (300 cells per well of a 12-well plate) on plates previously coated with 0.1% gelatin in N2B27+2i-LIF. After 5 days, the number of colonies was manually counted.

### Image analysis

All images were analyzed in Fiji/ImageJ (National Institutes of Health, Bethesda, MD, USA) (Schindelin et al., 2012). Raw images were used for quantification. The contrast was adjusted for clarity of presentation in the figures.

#### Abscission duration scoring in live cells

We scored abscission as the moment when we saw a rupture of microtubules in the bridge.

#### Number of bridges

The number of bridges was counted using the tubulin channel and the number of cells was counted using a nuclear marker (usually Aurora B or Hoechst). The number of bridges was then divided by the number of cells. A higher ratio of bridges/cells suggests that abscission is slow while low ratios suggest that abscission is fast.

#### Intensity of Aurora B/P-Aurora B/acetylated tubulin/total tubulin/ubiquitin-K48

To measure intensities in intercellular bridges on fixed samples, a Region Of Interest (ROI) was created surrounding the bridge using one of the tubulin channels. Then, the sum intensity projection was made of the slices in which a bridge was present. Only flat bridges (which was the majority of bridges) were selected for quantification. The mean Aurora B/P-Aurora B/tubulin/ubiquitin-K48 intensity was measured using the ROI. To calculate the mean intensity of Aurora B/P-Aurora B/tubulin/ubiquitin-K48 in the bridge, the background mean intensity was subtracted from the mean intensity of Aurora B/P-Aurora B/tubulin/ubiquitin K48 measured in the bridge.

### Statistical analysis and data display

In the graphs, the data is given as mean ± s.d., unless otherwise stated. Means of two groups were compared by an appropriate test (Student’s t-test if the data had normal distribution with a similar standard deviation, with Welch correction if the standard deviation was different, or Mann-Whitney if the distribution was not normal). The data shown in the images shows a representative intercellular bridge where similar observations were made in at least 10 bridges from at least 3 independent experiments. Different replicates are shown as variations of the main colors (for example, 3 shades of grey, orange or purple).

Graphs were generated in GraphPad Prism (MathWorks, Natick, MA, USA). Fiji was used to scale images and adjust brightness and contrast. Figures were assembled in Adobe Illustrator (Adobe Systems, Mountain View, CA, USA).

## Acknowledgements

We would like to thank all the members of the Chaigne lab and the whole department of Cell Biology, Neurobiology and Biophysics at Utrecht University for helpful discussions. We thank Anna Akhmanova and Benjamin P. Bouchet (Akhmanova lab, Utrecht University) for advice and discussions. We would like to thank Jesse Veenvliet (MPI Dresden) and Antoine Zalc (Institut Cochin) for helpful discussions and feedback.

This work was supported by Utrecht University and NWO (NWO-Vidi to A.C.).

## Author contributions

S.K. performed most of the experiments and analysis. B.M.L. performed the CLASP1 and CLASP2 fixed and live experiments and analysis. E.T. cloned some of the plasmids used in this study. A.C. supervised and conceptualized the experimental work and acquired funding. All authors discussed the data and the manuscript.

