## Supplementary material for "Wnt signalling controls abscission dynamics in mouse embryonic stem cells"

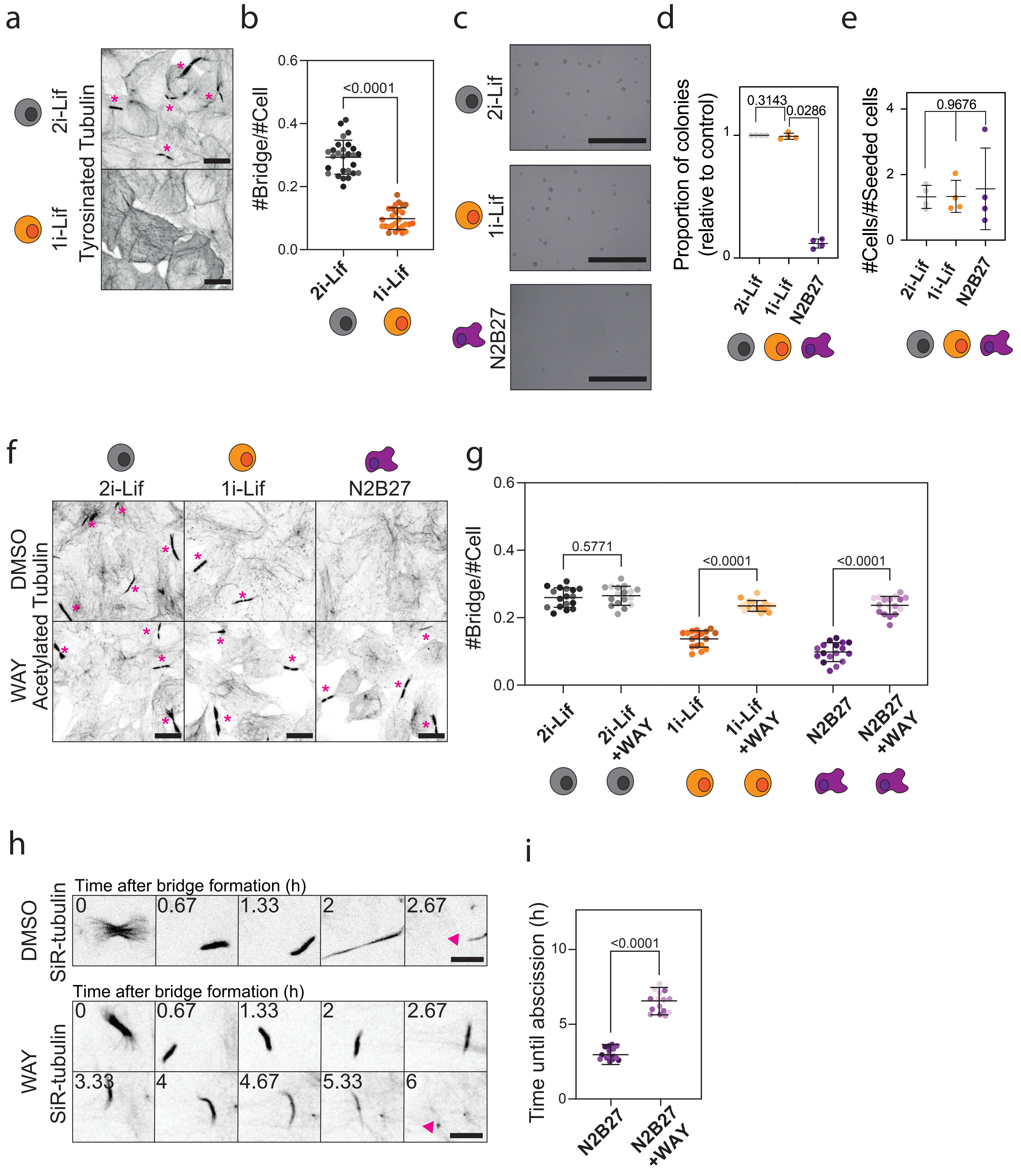


Supplementary figure 1

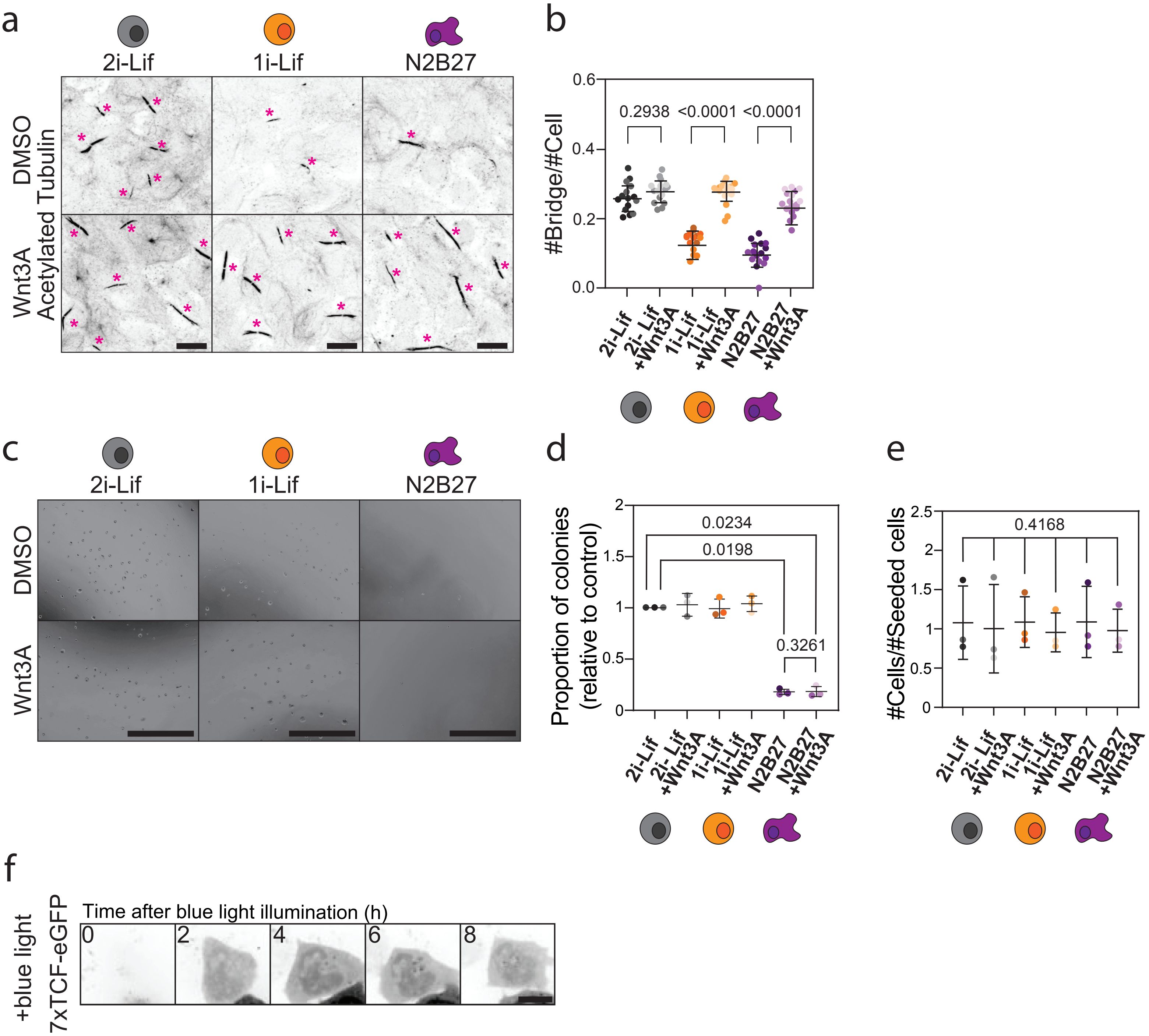


Supplementary figure 2

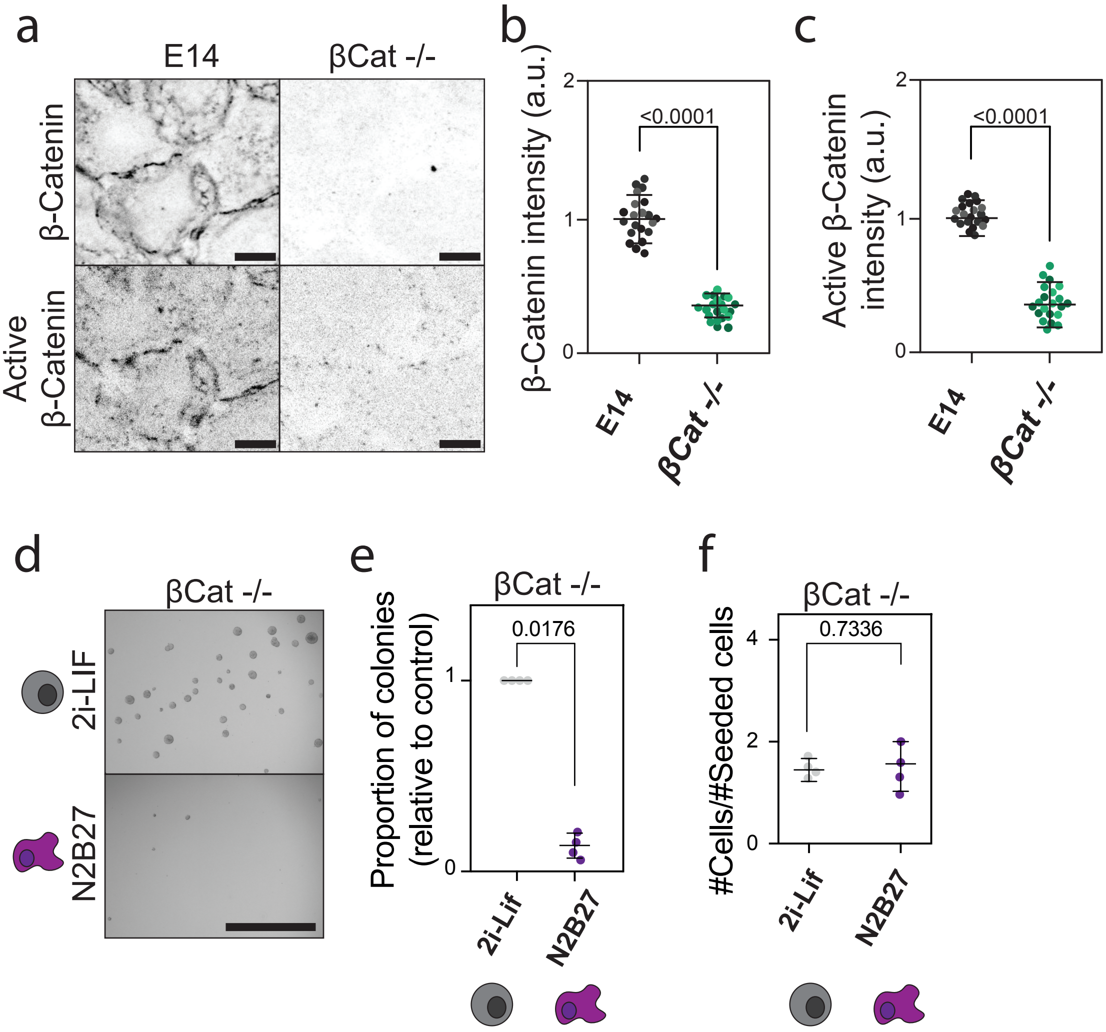


Supplementary figure 3

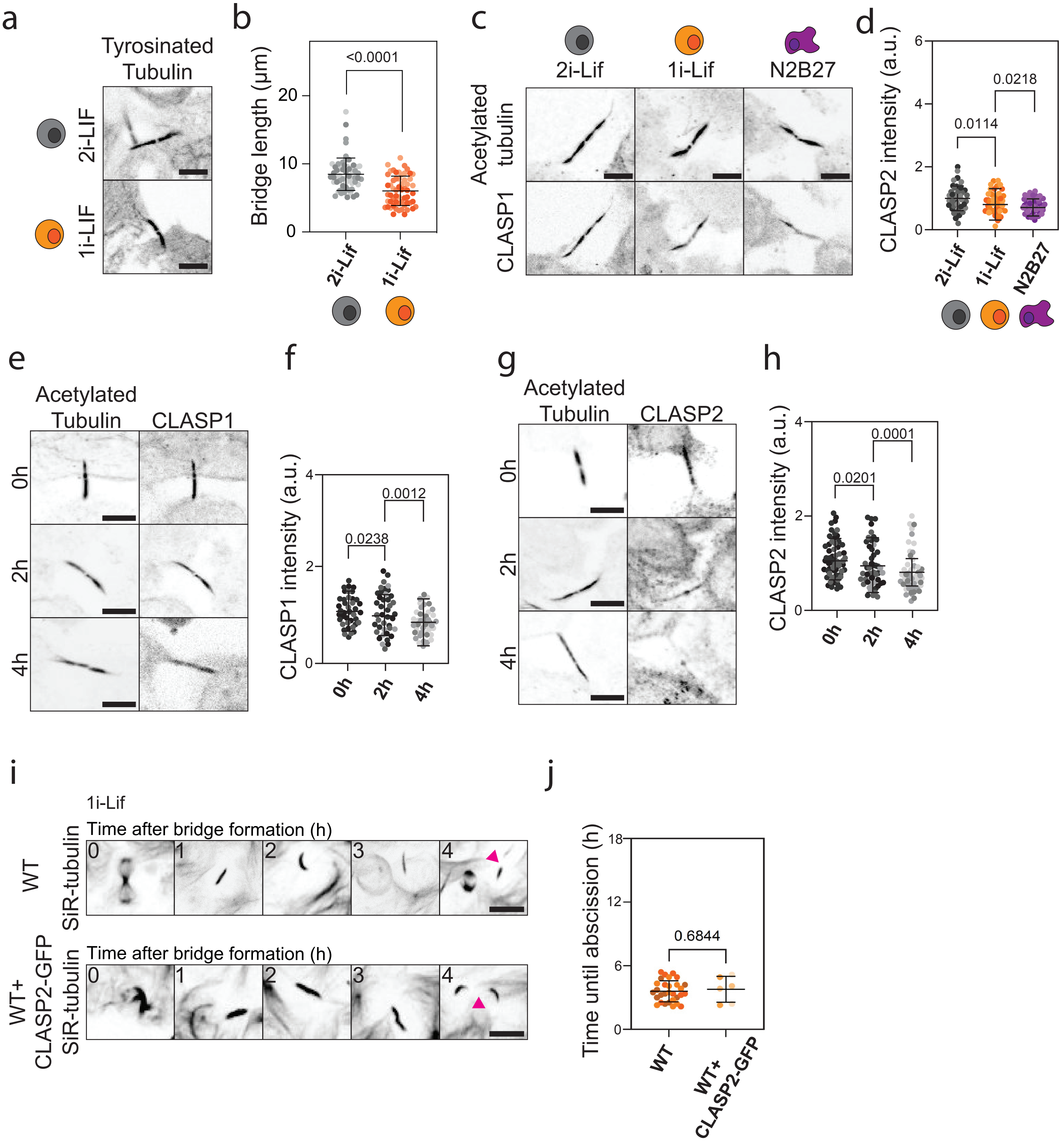


Supplementary figure 4

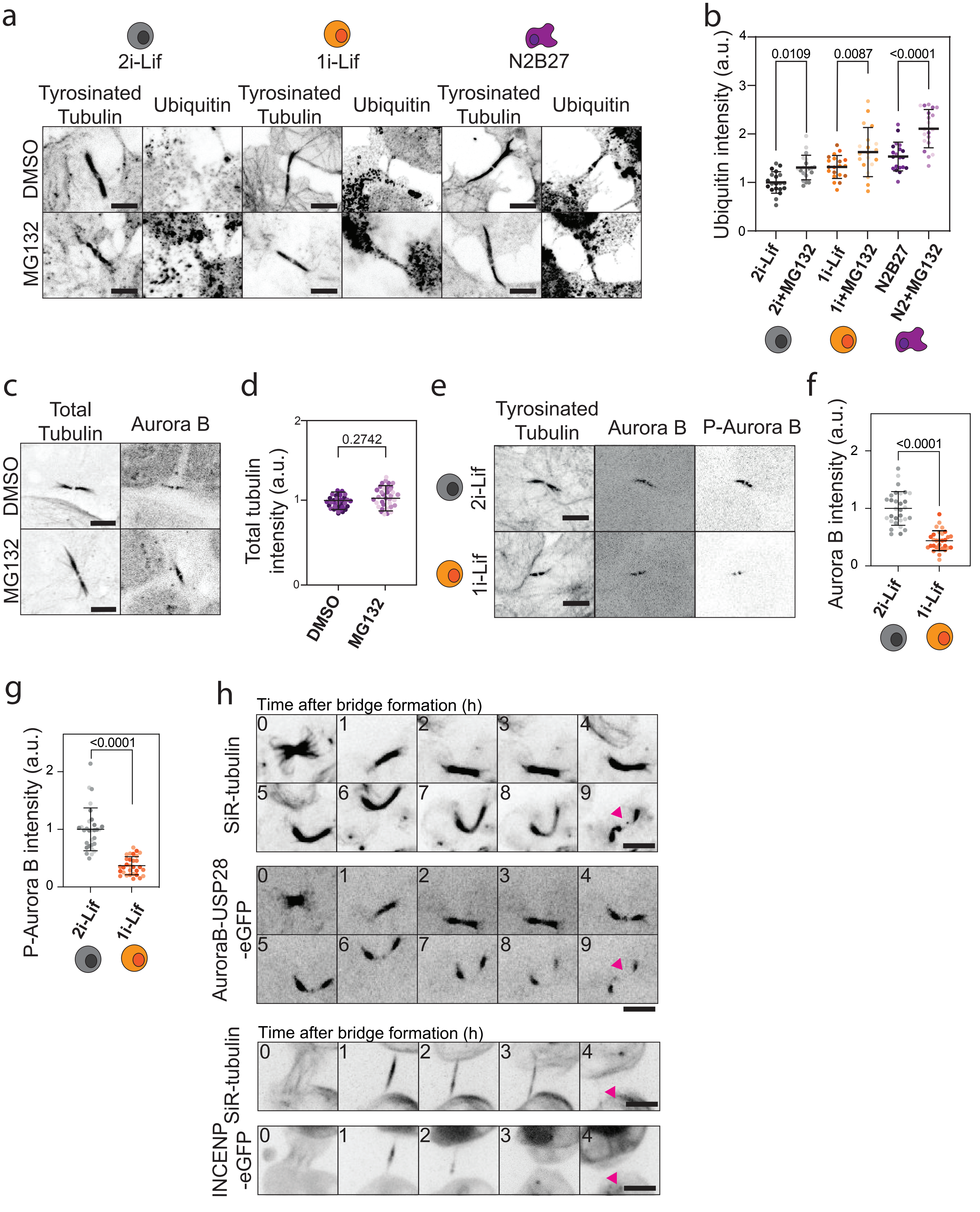


Supplementary figure 5

**Supplementary Figure 1. Abscission is delayed upon Wnt signalling activation in mouse embryonic stem cells.** a) Immunofluorescence showing the number of bridges in naïve ESCs in normal 2i-LIF (top) or 1i-LIF (bottom). A Z-projection over the height of the whole cell is shown. The bridges are shown with the staining of tyrosinated tubulin. Bridges are highlighted with pink asterisks. Scale bars: 10 μm. b) Quantification of the number of bridges per cell in naïve ESC in normal 2i-LIF (grey) or in 1i-LIF (orange). The mean and standard deviation are shown. N=3 replicates. c) Clonogenicity assay on mESCs grown in 2i-Lif (top), 1i-Lif (middle) and N2B27 (bottom) showing colonies formed after 20 h of respective media growth. Scale bars: 1 mm. d) Quantification of the number of colonies formed in 2i-Lif (grey), 1i-Lif (orange) and N2B27 (purple). The mean and standard deviation are shown. N = 4 replicates. e) Quantification of the number cell doublings in 2i-Lif (grey), 1i-Lif (orange) and N2B27 (purple). The mean and standard deviation are shown. N = 4 replicates. f) Immunofluorescence showing the number of bridges in mESCs in 2i-Lif (left), 1i-Lif (middle) and N2B27 (top right) and in the presence of Dkk1 inhibitor WAY-262611 (bottom row). A Z-projection over the height of the whole cell is shown. The bridges are shown with the staining of tyrosinated tubulin. Bridges are highlighted with pink asterisks. Scale bars: 10 μm. g) Quantification of the number of bridges per cell in mESC in 2i-Lif (dark grey), 1i-Lif (dark orange) and N2B27 (dark purple) treated with DMSO or 2i-Lif (light grey), 1i-Lif (light orange) and N2B27 (light purple) treated with WAY-262611. The mean and standard deviation are shown. N=3 replicates. h) Live-cell imaging of DMSO (top) and WAY-262611 (bottom) treated mESCs in N2B27 media and incubated overnight with 20 nM SiR-tubulin. A Z-projection over the height of the whole cell is shown. Tubulin is shown in black. The pink arrows indicate the cut sites. One frame is shown every 0.5h. Scale bars: 10 μm. i) Quantification of the duration of abscission from bridge formation until microtubule severing in mESCs treated with DMSO (dark purple) and WAY-2626211 (light purple) in N2B27. The mean and standard deviation are shown. N=3 replicates.

**Supplementary Figure 2. Activation of Wnt signalling delays abscission in mouse embryonic stem cells exiting naïve pluripotency.** a) Immunofluorescence showing the number of bridges in mESCs in 2i-Lif (top left), 1i-Lif (top middle) and N2B27 (top right) in the presence of Wnt3A ligand (bottom row). A Z-projection over the height of the whole cell is shown. The bridges are shown with the staining of acetylated tubulin. Bridges are highlighted with pink asterisks. Scale bars: 10 μm. b) Quantification of the number of bridges per cell in mESC in 2i-Lif (dark grey), 1i-Lif (dark orange) and N2B27 (dark purple) or 2i-Lif (light grey), 1i-Lif (light orange) and N2B27 (light purple) with Wnt3A ligand. The mean and standard deviation are shown. N=3 replicates. c) Clonogenicity assay of mESCs treated with Wnt3A ligand showing colonies formed after 20 h in 2i-Lif (top left), 1i-Lif (top middle), N2B27 (top right), 2i-Lif + Wnt3A (bottom left), 1i-Lif + Wnt3A (bottom middle) and N2B27+Wnt3A (bottom right). Scale bars: 1 mm. d) Quantification of the number of colonies formed in 2i-Lif (dark grey), 2i-Lif+Wnt3A (light grey), 1i-Lif (dark orange), 1i-Lif+Wnt3A (light orange), N2B27 (dark purple) and N2B27+Wnt3A (light purple). The mean and standard deviation are shown. N = 4 replicates. e) Quantification of the cell proliferation in 2i-Lif (dark grey), 2i-Lif+Wnt3A (light grey), 1i-Lif (dark orange), 1i-Lif+Wnt3A (light orange), N2B27 (dark purple) and N2B27+Wnt3A (light purple). The mean and standard deviation are shown. N = 4 replicates. f) Live-cell imaging of mESCs in N2B27 media transfected with 7xTCF-GFP. Blue light illumination started at the beginning of imaging. A Z-projection over the height of the whole cell is shown. One frame is shown every 2h. Scale bar: 10 μm.

**Supplementary Figure 3. Canonical Wnt signalling does not influence abscission dynamics.** a) Immunofluorescence showing the localization of β-catenin (top) and active β-catenin (bottom) at the bridge in WT mESCs (left) or β-catenin KO line (right) in 2i-Lif. A Z-projection over the height of the whole sample is shown. The bridges are shown with the staining of acetylated tubulin. Scale bars: 10 μm. b) Quantification of β-catenin intensity in WT mESCs (grey) and β-catenin KO line (green) in 2i-Lif. The mean and standard deviation are shown. N=3 replicates. c) Quantification of active β-catenin intensity in WT mESCs (grey) and β-catenin KO line (green) in 2i-Lif. The mean and standard deviation are shown. N=3 replicates. d) Clonogenicity assay on β-catenin KO line showing colonies formed after 20 h of naïve pluripotency (top) or exit from naïve pluripotency (bottom). Scale bars: 1 mm. e) Quantification of the number of colonies formed in 2i-Lif (grey) and N2B27 (purple) in β-catenin KO line. The mean and standard deviation are shown. N = 4 replicates. f) Quantification of the number cell doublings in 2i-Lif (grey) and N2B27 (purple) in β-catenin KO line. The mean and standard deviation are shown. N = 4 replicates.

**Supplementary Figure 4. Wnt signalling increases microtubule acetylation at the bridge.** a) Immunofluorescence showing the localization of tyrosinated tubulin at the bridge in 2i-Lif (top) and 1i-Lif (bottom). Scale bar: 5 μm. b) Quantification of the bridge length in mESCs in 2i-Lif (grey) and 1i-Lif (orange) media. The mean and standard deviation are shown. N=3 replicates. c) Immunofluorescence showing the localization of CLASP1 at the bridge in 2i-Lif, 1i-Lif and N2B27(bottom row, from left to right, respectively). A Z-projection over the height of the whole sample is shown. The bridges are shown with the staining of acetylated tubulin. Scale bar: 5 μm. d) Quantification of CLASP1 intensity in mESCs in 2i-Lif (grey), 1i-Lif (orange) and N2B27 (purple). The mean and standard deviation are shown. N=3 replicates. e) Immunofluorescence showing the localization of CLASP1 in mESCs in 2i-Lif during bridge maturation at bridge formation (0h), 2 h after bridge formation (2h), and 4 h after bridge formation (4h). Scale bars: 5 µm. f) Quantification of CLASP1 intensity in mESCs in 2i-Lif media during bridge maturation at 0h (dark grey), 2h (grey) and 4h (light grey) after bridge formation. The mean and standard deviation are shown. N=3 replicates. g) Immunofluorescence showing the localization of CLASP2 in mESCs in 2i-Lif during bridge maturation at bridge formation (0h), 2 h after bridge formation (2h), and 4 h after bridge formation (4h). Scale bars: 5 µm. h) Quantification of CLASP2 intensity in mESCs in 2i-Lif media during bridge maturation at 0h (dark grey), 2h (grey) and 4h (light grey) after bridge formation. The mean and standard deviation are shown. N=3 replicates. i) Live-cell imaging of WT (top) and WT+CLASP2-GFP (bottom) incubated overnight with 20 nM SiR-tubulin in 1i-Lif . A Z-projection over the height of the whole cell is shown. Tubulin is shown in black. The pink arrows indicate the cut sites. One frame is shown every 1h. Scale bars: 10 μm. j) Quantification of the duration of abscission from bridge formation until microtubule severing in mESCs in WT (dark grey) and WT+CLASP2-GFP (light grey) mESCs in 1i-Lif. The mean and standard deviation are shown. N=3 replicates.

**Supplementary Figure 5. Protein degradation leads to faster abscission.** a) Immunofluorescence showing the localization of tyrosinated tubulin (left) and ubiquitin (right) at the bridge in mESCs in 2i-Lif (left), 1i-Lif (middle) or N2B27 (right) treated with DMSO (top) or MG132 (bottom). A Z-projection over the height of the whole sample is shown. The bridges are shown with the staining of tyrosinated tubulin. Scale bars: 5 μm. b) Quantification of ubiquitin intensity in mESCs in 2i-Lif (dark grey), 2i-Lif+MG132 (light grey), 1i-Lif (dark orange), 1i-Lif+MG132 (light orange), N2B27 (dark purple) and N2B27+MG132 (light purple). The mean and standard deviation are shown. N=3 replicates. c) Immunofluorescence showing the localization of total tubulin (left) and Aurora B (right) at the bridge in mESCs in N2B27 treated with DMSO (top) or MG132 (bottom). A Z-projection over the height of the whole sample is shown. Scale bar: 5 μm. d) Quantification of the intensity of total tubulin in mESCs in N2B27 treated with DMSO (dark purple) or MG132 (light purple). The mean and standard deviation are shown. N=3 replicates. e) Immunofluorescence showing the localization of tyrosinated tubulin (left), Aurora B (middle) and P-Aurora B (right) at the bridge in 2i-Lif (top) and 1i-Lif (bottom). A Z-projection over the height of the whole sample is shown. The bridges are shown with the staining of tyrosinated tubulin. Scale bar: 10 μm. f) Quantification of Aurora B intensity in mESCs in 2i-Lif (grey) and 1i-Lif (orange). The mean and standard deviation are shown. N=3 replicates. g) Quantification of P-Aurora B intensity in mESCs in 2i-Lif (grey) and 1i-Lif (orange). The mean and standard deviation are shown. N=3 replicates. h) Live-cell imaging of mESCs transfected with Aurora B-USP28-eGFP (second row) and added 20 nM SiR-tubulin (first row), and INCENP-eGFP (fourth row) with added 20 nm SiR-tubulin (third row) in N2B27. The cell transfected with Aurora B-USP28-eGFP is the same cell as in Figure 5l. A Z-projection over the height of the whole cell is shown. The pink arrows indicate the cut sites. One frame is shown every 1h. Scale bars: 10 μm.

**Movie legend**

**Movie 1**: Live-cell imaging of mESCs in 2i-LIF (left) and 1i-LIF (right) incubated overnight with 20 nM SiR-tubulin. A Z-projection over the height of the whole cell is shown. Tubulin is shown in black. One frame every 5 min. Scale bars: 10 μm.

**Movie 2**: Live-cell imaging of mESCs in N2B27 with DMSO (left) or Wnt3A ligand (right) incubated overnight with 20 nM SiR-tubulin. A Z-projection over the height of the whole cell is shown. Tubulin is shown in black. One frame every 5 min. Scale bars: 10 μm.

**Movie 3**: Live-cell imaging of mESCs transfected with Tubulin-GFP (left) or Tubulin-GFP and Opto-Wnt (LRP6-Cry-mCherry, right). A Z-projection over the height of the whole cell is shown. Tubulin is shown in black. One frame every 5 min. Scale bars: 10 μm.

**Movie 4**: Live-cell imaging of WT (left) and β-catenin KO mESCs (right) incubated overnight with 20 nM SiR-tubulin. A Z-projection over the height of the whole cell is shown. Tubulin is shown in black. One frame every 5 min. Scale bars: 10 μm.

**Movie 5**: Live-cell imaging of mESCs in 2i-Lif incubated overnight with 20 nM SiR-tubulin (left) and transfected with GSK-3β-GFP (right). A Z-projection over the height of the whole cell is shown. One frame every 5 min. Scale bar: 5 μm.

**Movie 6**: Live-cell imaging of WT (left) and WT+CLASP2-GFP (right) incubated overnight with 20 nM SiR-tubulin in 2i-Lif. A Z-projection over the height of the whole cell is shown. Tubulin is shown in black. One frame every 5 min. Scale bars: 10 μm.

**Movie 7**: Live-cell imaging of WT (left) and WT+AuroraB-USP28-eGFP (right) incubated overnight with 20 nM SiR-tubulin in N2B27. A Z-projection over the height of the whole cell is shown. Tubulin is shown in black. One frame every 5 min. Scale bars: 10 μm.
